# Optimum Push-off During Uneven Walking for Just-in-Time Strategy; Delayed Push-off Exertion is Mechanically Costly

**DOI:** 10.1101/2024.07.14.603455

**Authors:** Seyed Saleh Hosseini-Yazdi

## Abstract

It is shown that step mechanical work roughly describes walking energetics, and optimal walking economy is achieved by pre-emptive step work. We suggest this is also true for uneven walking. Using a simple powered walking model, we estimated the preferred pre-emptive push-offs to cover the entire step energy. The maximum push-off is exerted when the subsequent heel-strike dissipation is zero, setting an upper bound for step-up amplitude achievable with pre-emptive push-off. For instance, at a walking speed of 1.4 m · s^−1^, the maximum step-up is 0.106 m. Conversely, for any step-up amplitude, there is a minimum walking speed. For a step-up height (Δh) of 0.06 m, the minimum walking speed is 1.06 m · s^−1^. The importance of pre-emptive push-off and optimal timing of push-off and collision is widely discussed. However, there are cases where this timing is undermined, such as during uneven walking, necessitating post-transition mechanical energy compensation. The ankle (via delayed push-off) or hip can provide mid-flight energy, but no mechanical determinant prefers one source over the other. Our modeling demonstrates that delayed push-off entails mechanical energy waste, likely converted to heat by stretching the stance leg. This stretch may also release energy stored during the heel-strike (e.g., in the Achilles tendon), exacerbating the required mechanical work performance in the subsequent step transition. Hence, we propose that during the double support phase, when the stance leg is switched, hip actuation becomes mechanically preferable. Physiological observations also support our proposition.

## Introduction

Minimizing energy expenditure is one of the major determinants of human walking characteristics [1,2]. As such, it is proposed that humans select a combination of walking variables that coincide with optimum walking energetics [1]. Such features are termed as nominal walking parameters [3,4]. Any deviation from nominal gait is suggested to associate with elevated metabolic rates [4]. Since it is suggested that the best walking economy is achieved by pre-emptive push-off [3], any delayed push-off or other forms of positive energy inducement increases the mechanical cost of walking [1]. Instances such as uneven walking may enforce delayed energy inducements [5]. Any post transition or mid-flight mechanical energy addition must also be mechanically efficient. The ankle and hip are nominated as the energy inducement sources, yet their work performance ramifications are not evaluated.

It is hypothesized that the pre-emptive push-off limits the subsequent collision work magnitude [3,6]. The optimum push-off (minimum) [5,7] equates to the negative collision work to maintain a steady walking velocity (Figure *1*A) [3,8]. Therefore, during the stance phase, no further energy inducement is necessary [6]. For nominal walking Donelan et al. [9] suggestion that the positive work exerted during the mid-flight (rebound) phase is the release of stored energy in tendons and tissues, and it is completely absorbed during the preload, support this hypothesis. Therefore, it does not require active mechanical work generation that incurs metabolic expenditure [1,10].

**Figure 1:**
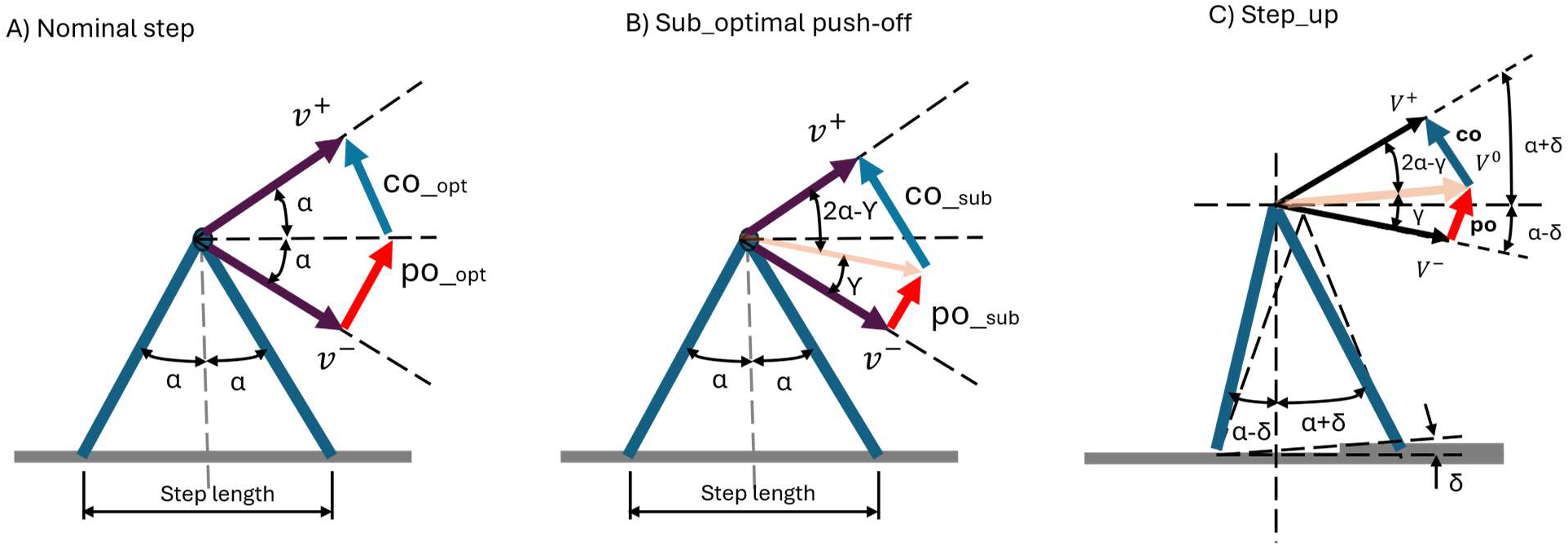
The step-to-step transition is when the stance leg is switched. (A) During a nominal walking, the push-off work is exerted pre-emptively and is equal to the subsequent collision. (B) If a sub-optimal push-off is exerted, the subsequent collision magnitude grows. (C) Uneven walking is a deviation from nominal even walking. In ideal case, the pre-emptive push-off must provide the mechanical energy for step change and gravity work.

If the relative timing of push-off and collision is undermined, the randomly exerted pre-emptive push-off may not completely adjust the COM velocity mid step-to-step transitions that yields to heel strike collision growth [3,11,12]. Therefore, at post transition, further positive mechanical work (muscle work) is required to sustain the gait (Figure *1*B). Such work that is performed mid-flight is termed as rebound [6]. If the rebound work is used to energize the progression (exceeding the preload), it becomes an active work that costs metabolic energy similar to step transition work [9].

During the uneven walking, the push-off not only has to cancel the collision dissipation, but also provides the energy required to go atop bumps (Figure *1*C) [13]. Hence, it is conceivable to observe larger push-offs and smaller collisions during step-ups [5,13]. The difference exacts the work against gravity that is converted to gravitational potential energy [13]. Nevertheless, one consequence of uneven walking might be upsetting the optimal timing of push-off and collision [5]. Therefore, like some even walking instances, the entire necessary mechanical energy might not be provided pre-emptively.

Uneven walking requires active regulation to maintain a sustained gait [5,13]. If terrains’ complexity intensities allow humans to plan for the extended horizon [14], humans demonstrate a tri-phasic control before and after any particular complexity encounter [15]. On the other hand, it is also proposed that extending the regulation over several steps is adding to the estimates’ errors [16]. Darici and Kuo [14] have also reported that the advantage of forward lookahead horizon saturates beyond six to eight steps. Therefore, it is proposed that humans tend to utilize the Just-In-Time application of the visual information over complicated terrains [16,17]. Since the just-in-time regulation relies on each step’s performance, the walkers may suffer a higher cost of mechanical work performance than the anticipatory approach [14,15]. The brief background provided demonstrates that almost all of the walking simulations and experimental works are focused on the impact of pre-emptive push-off and consequence of sub-optimal push-off and collision exertions. Here, we utilize a simple powered walking model to estimate the required pre-emptive (preferred) push-off that energizes the entire step-up for a given walking speed. We also estimate the associate bounds based on step-up amplitude increase. Since there is no mechanical comparison between the sources of the post transition energy, we additionally evaluate the costs of work performances by the prior trailing ankle versus planted leading leg hip actuation when the stance leg is switched.

## Simulations

### Step-up with a pre-emptive push-off

Without losing the generality, we consider a step-up to describe the consequences of uneven walking [15] and the associated deviation from the even walking step work (push-off). We use a simple walking model [3] to evaluate the optimum step work (push-off) that provides the entire energy of the step-to-step transition and the work against gravity. For a given walking speed (*ν*_*ave*_), there is an optimum step length that coincides with the minimum energy expenditure [1,2]. For the small angle approximation, the step length may be represented by the angle between the leading and trailing leg at the point of transition (double support phase). It is indicated by 2*α* [3].

At the end of a stance phase and beginning of the double support, we show the COM velocity by *ν*^−^, the velocity in the middle of transition by *ν*^0^, and the resulted velocity after the transition by *ν*^+^. The *ν*^−^ is approximately equal to *ν*_*ave*_ [3]. We may derive the optimal push-off work by two method which are essentially similar (Figure *1*C):

#### Net work method

the post transition net mechanical work must be equal to the step-up gravity work (*PO* − *CO* = *g* · Δ*h*, in which Δ*h* represents step-up elevation change). For a random exerted push-off (*po*), the mid-transition COM velocity angle with the pre-transition velocity is 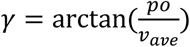, thus, subsequent dissipation impulse becomes (Figure *2*A):

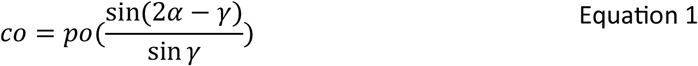

**Figure 2:**
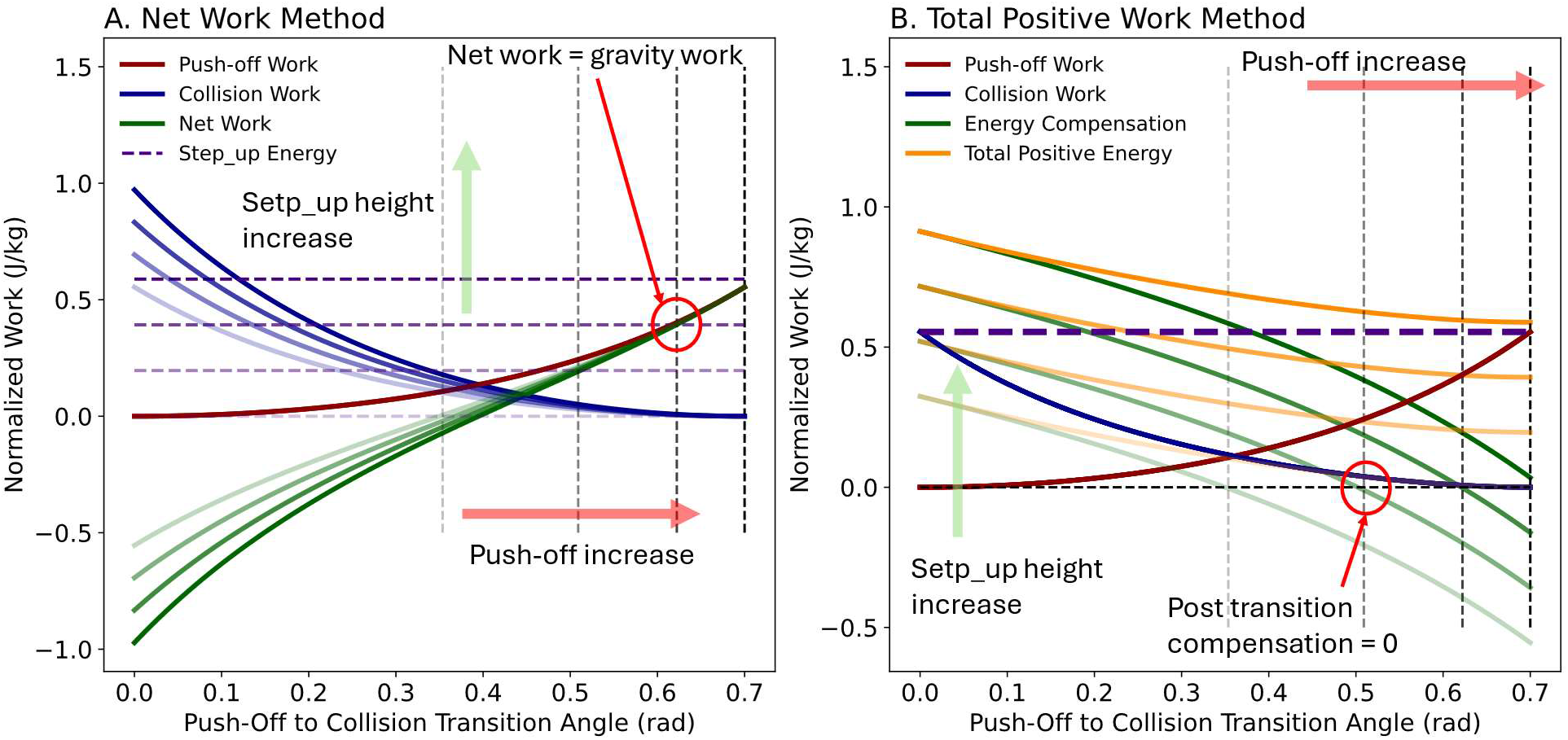
To calculate the optimum push-off work that covers the step-to-step transition and gravity works, two methods are presented: (A) net work method: the post transition work (push-off – collision) must be equal to the gravity work. (B) The post transition compensation must be zero when before and after transition average speeds are the same. The analysis was performed for v = 1.25 *m* · *s*^−1^ (α = 0.35 rad.) as the step-up amplitude ranged from 0 m to 0.06 m. Push-off to collision transition angle represents the COM velocity mid-transition with pre-transition COM velocity.

Calculating the mechanical work associated with push-off and collision impulses as 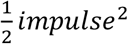 [5], the optimum push-off is: *PO* = *CO* + *g* · Δ*h*, therefore:

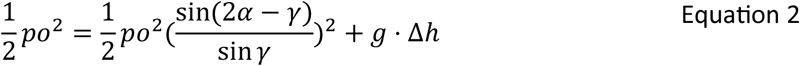

Rearranging the equation:

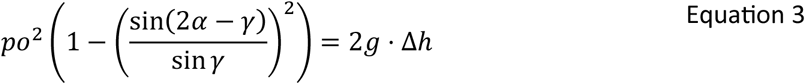

As such, the optimum push-off must satisfy Equation 3.

#### Post transition compensation method

in this method, the post transition kinetic energy must sustain the average walking velocity (*ν*_*ave*_) and cover the step-up gravity work. Therefore, the post transition (mid-flight) mechanical energy compensation must be zero. Therefore (Figure *2*B):

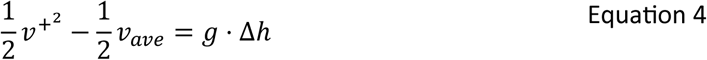

As such, the optimum post transition velocity must satisfy the following equation:

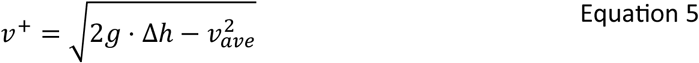

Since for a given push-off, the post transition velocity is:

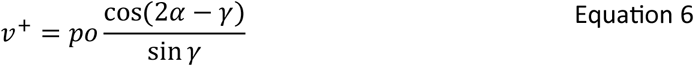

As such, the optimum push-off must satisfy the

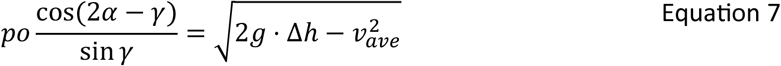

### Post step transition mechanical work compensation

We assumed that step transition mechanical work is performed in two separate phases while both legs are in contact with the ground (double support). In the first phase, the COM rotates about the trailing leg and the mechanical work is performed pre-emptively along the trailing leg (Figure 3A). “*The post-transition work*” compensation is necessary if the pre-emptive step work does not provide all the required energy. For even walking, the post transition work is ′*Collision* − *Push*_*off*′, whereas for a step-up it is ′*Collision* + *g* · Δ*h* − *push*_*off*′. In post transition, the leading leg becomes the COM rotation base.

**Figure 3:**
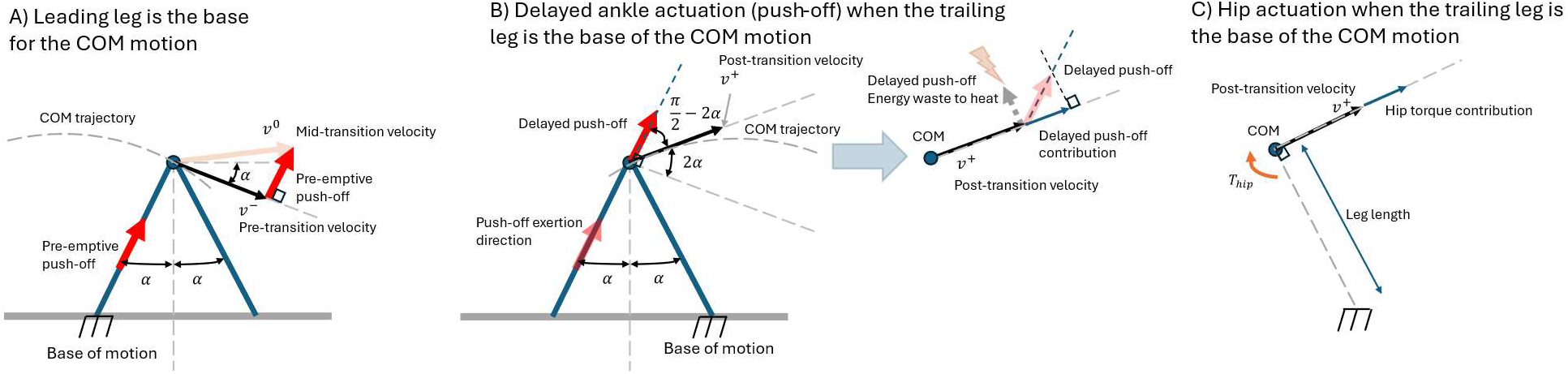
(A) during the step-to-step transition, the pre-emptive push-off partially redirects the COM velocity. The subsequent collision adjusts COM velocity completely for the next stance. (B) After the stance leg switch, if the trailing leg exerts any further push-off work (delayed push-off), since push-off impulse and COM velocity trajectories differ, a portion of induced push-off is wasted. (C) If the required post transition mechanical energy is covered by the stance leg hip actuation, it provides an angular impulse that its work is equal to the mechanical energy deficit.

If the remainder of the post transition compensation comes from the trailing leg ankle, its impulse still will be induced along the trailing leg. As such, the portion that contributes to the COM velocity becomes (contribution impulse or ‘*cont*’, Figure 3B):

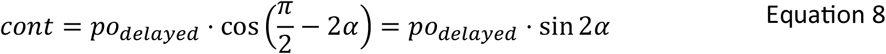

Therefore, when its mechanical energy contribution is equal to the needed compensation(‘*COMP*’):

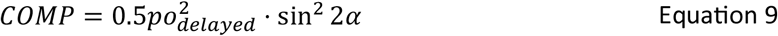

While the effective (true) trailing leg’s ankle work performance as 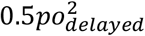, hence:

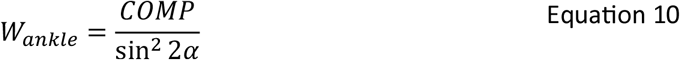

On the other hand, if hip covers the post transition work, its actuation exerts a torque (angular impulse) to the COM. Hence, its work becomes equal to the post transition work (*W*_*hip*_ *= COMP*, Figure 3C).

## Results

We utilized a powered simplest walking model [3] to derive the analytical optimum pre-emptive push-offs. For any given speed, there was a minimum optimal push-off that increased with the speed magnitude. It indicated the optimal push-off for even walking to sustain the kinetic energy after the step transition. The push-off trajectories associated with each velocity exhibited a maximum step-up amplitude. The walking speed rise increased the step-up amplitude attainable only if the pre-emptive push-off would be used. For instance, for v = 1.4 m · s^−1^, simulation indicated that the walker could go atop a step-up with maximum amplitude of Δh = 0.106 m if the pre-emptive push-off provided the entire step work. We also observed that for Δh = 0.04 m and 0.06 m, there were minimum walking speed distinguished (v = 0.86 m · s^−1^ and 1.06 m · s^−1^, respectively, Figure 4) in the speed range examined

**Figure 4:**
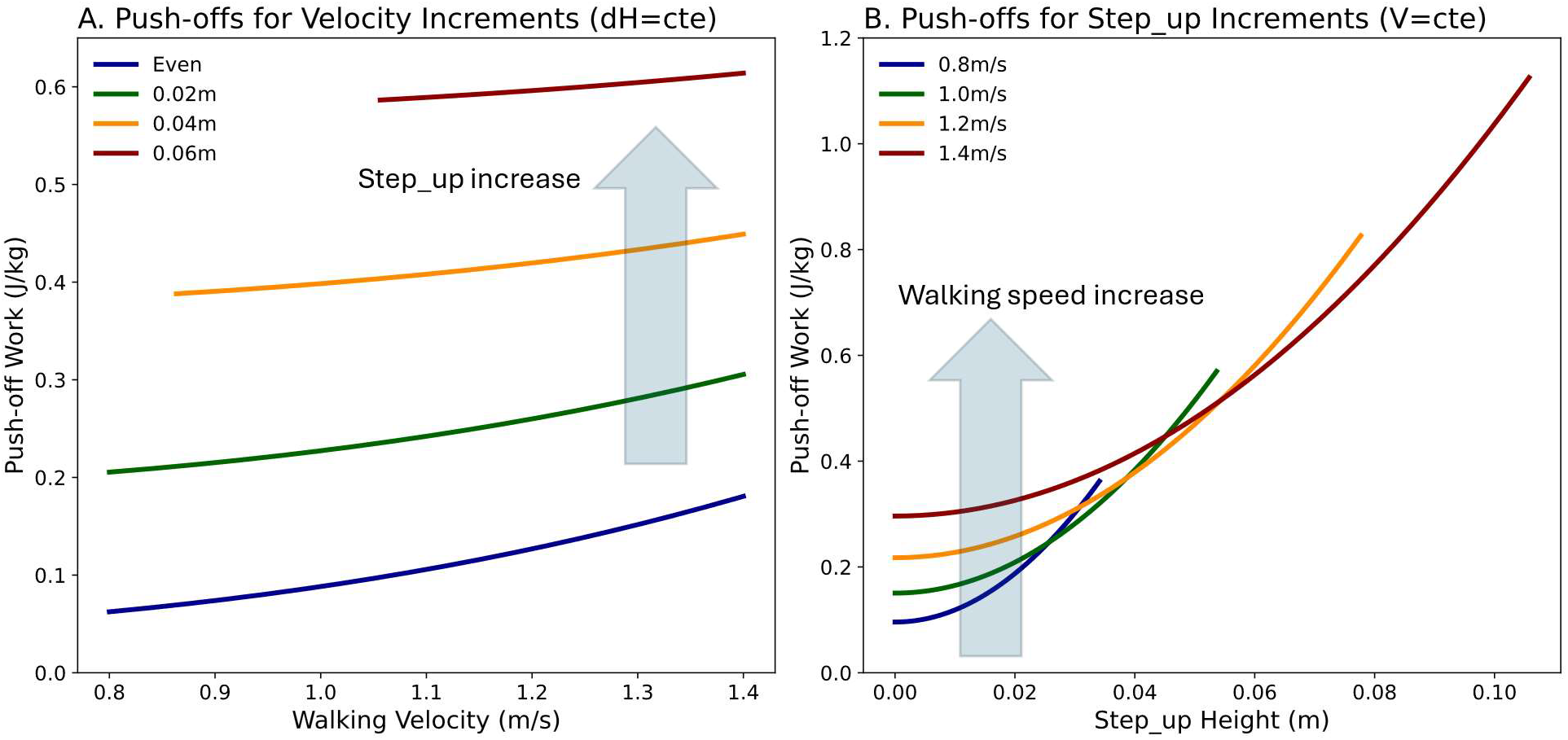
(A) for any step-up, the average walking speed has a lower bound in which the entire gait positive mechanical work is provided by the pre-emptive push-off. (B) For any given walking speed, there is an upper bound for the step-up magnitude in which the entire mechanical energy is provided pre-emptively.

For even walking, the pre-emptive push-off was suboptimal (v = 1.25 m · s^−1^) was associated with increased collision impulse and its associated work. As such, the post transition compensation grew when the pre-emptive push-off declined. With optimal push-off exertion, collision and push-off were equal (0.46 m · s^−1^, or 0.1 J · kg^−1^) and the post transition compensation was zero. If the collision superseded the push-off, the collision grew to 3.2 time of the optimal push-off magnitude and the post transition compensation was equal to the collision (Figure 5).

**Figure 5:**
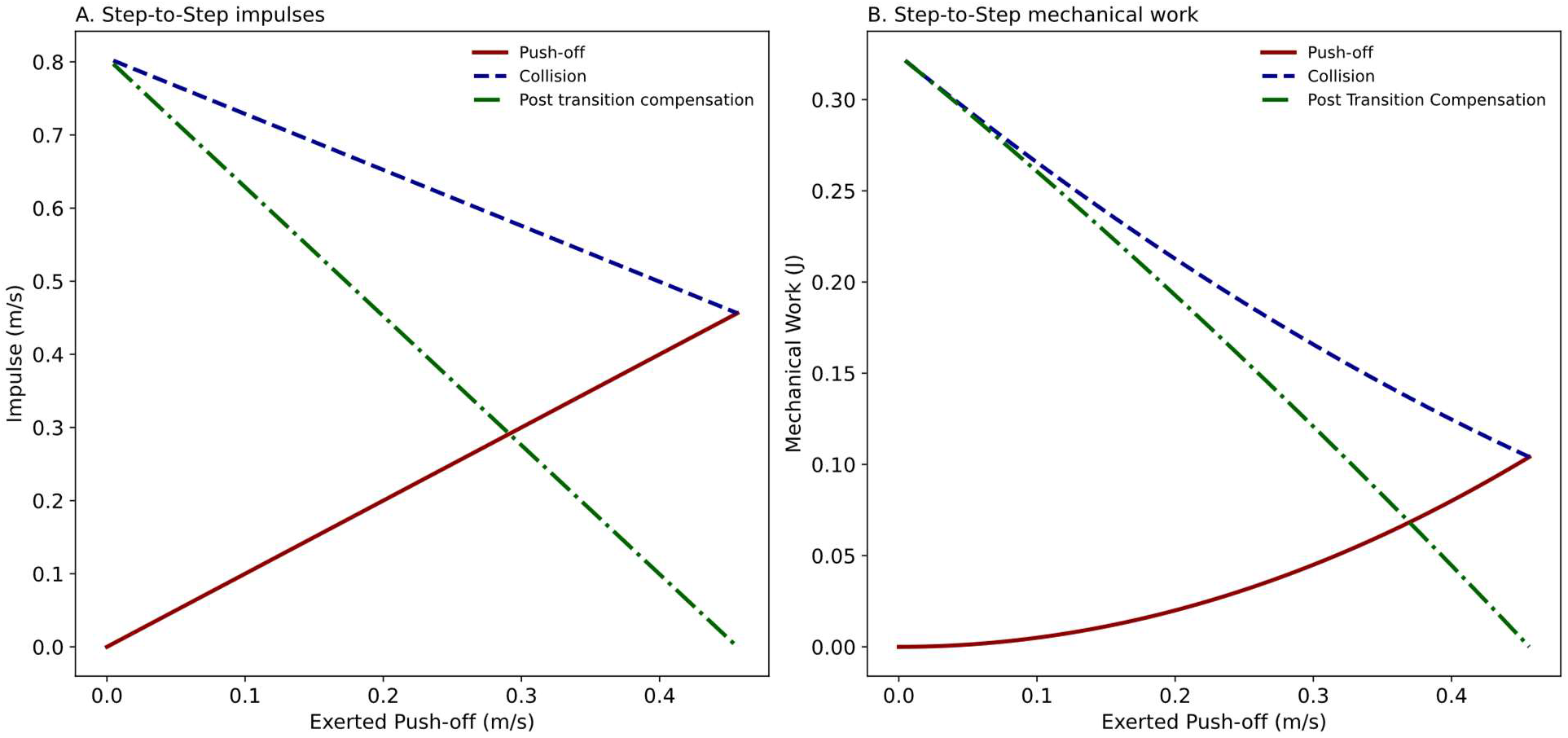
(A) when a sub-optimal push-off impulse is exerted, the subsequent collision impulse magnitude grows. Their difference declines as the induced push off approaches the nominal value (v = 1.25 m · s^−1^, even walking and α = 0.35 rad.). (B) Similar to impulses, with a sub-optimal push-off work, the magnitude of collision increases. The difference of push-off and collision work reduces as the push-off work gets closer to its nominal magnitude. At optimal push-off, the difference is zero. The sub-optimal push-off and collision difference must be compensated after the step transition during the subsequent mid-flight phase.

When the hip actuation provided the post transition compensation, its work was equal to the required compensation. On the contrary, the ankle actuation resulted in elevated mechanical cost. For even walking, the completely delayed push-off work was 7.5 times of the optimal push-off work (v = 1.25 m · s^−1^), whereas for Δh = 0.025 m, the completely delayed push-off mechanical was 5.0 time of the optimal push-off. The similar hip works were 3.2 and 2.1 times of the optimal push-off, respectively (Figure 6).

**Figure 6:**
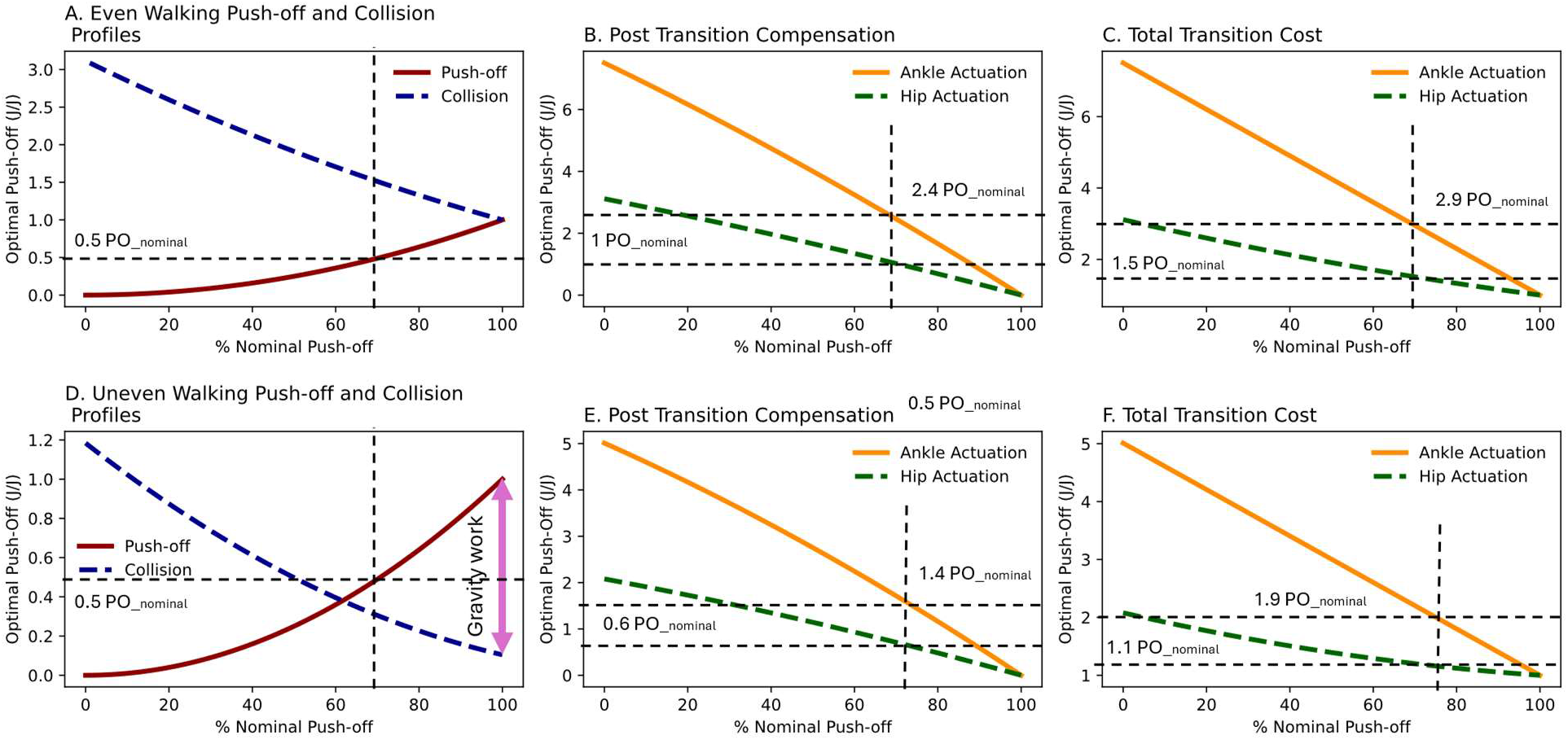
If sub-optimal push-off is exerted, the sub-optimal push-off and its subsequent collision difference may be compensated by a delayed push-off or hip actuation. For even (A. to C., v = 1.25 m · s^−1^) and uneven walking (D. to F., Δh = 0.025 m), the mechanical cost of delayed ankle actuation is much higher than hip actuation. The horizontal axis represents the pre-emptive push-off as a fraction of optimal push-off impulse (% of v · v^−1^), and the vertical axis represented the fraction of the work performance according to the optimal push-off (J · J^−1^). For even walking (top row), when pre-emptive push-off was 70% of optimal push-off impulse, the post transition hip compensation work was 1.0 of optimal push-off work, and the total step work reached 1.5 times of the optimal push-off work. If the compensation was provided by delayed push-off instead (ankle), the post transition compensation was 2.4 times of the optimal push-off and the total step work climbed to 2.9 times of the optimal push-off work. For uneven walking (lower row), the pre-emptive push-off equal to 70% of the optimal push-off required 0.6 and 1.4 times of the optimal push-off work if compensated by the hip and ankle actuations, respectively. Subsequently, the associated total step works were 1.1 and 1.9 times of the optimal push-off work.

## Discussion

We demonstrated two methods based on maintenance of the kinetic energy to estimate the preferred (pre-emptive) push-off for a given step-up, knowing the initial walking speed. Whether using the net work to overcome a particular step-up or reducing the post transition mechanical energy compensation to zero, we derive the same result for the optimal push-off. However, there are instances that during human walking, the proper order of push-off and collision is undermined and as such, mid-flight extra mechanical work addition becomes necessary. The main question becomes, is there any determinant to select a preferred method to energize the gait post transition. Since it appears that there is no mechanical determinant presented for this purpose, we investigated the ankle and hip joints impulse exertions during the double support phase when that stance phase is switched.

The pre-emptive push-off is bounded by the speed and step-up elevation. The pre-emptive push-off stems from the walker’s kinetic energy [3]. It is shown that for a given walking speed, there is a maximum performable push-off work that reduces the subsequent collision to zero [1,3]. If the maximum push-off cannot cover the needed gravity work, the remainder has to be made up for during the following stance phase [6]. As such, it defines an upper bound for the step-up amplitude for a given speed. Similar restriction may be expressed for an assumed step-up amplitude for which the maximum push-off defines a lower bound for the walking speed.

The importance of the optimal push-off and collision timing [11], or the preferred order of push-off and collision is discussed immensely [5,18]. We also demonstrated that with the pre-emptive push-off decline (random push-off as a fraction of optimum push-off), the subsequent collision grows exponentially. For instance, if the push-off is completely exerted after the heel strike, the collision magnitude is maximized [3]. The same is also true for the uneven walking (e.g. step-up). The only difference is the further collision reductions as greater push-offs are performed [13].

The source of the post transition mechanical works is widely neglected. The necessity of post transition mechanical work compensation when sup-optimal push-off is applied, is agreed [6]. Nevertheless, it is neglected if the source of the compensation has any mechanical implications. Therefore, simulations usually are indifferent about it [5,13]. Hip and ankle are mentioned as two possible sources for midstance energy inducements [3,9]. Based on the vertical ground reaction force or COM power trajectories, it is observed that a portion of push-off is exerted with delay, even for the nominal walking [19]. Yet, the heightened proximal muscle activities during uneven walking [20] might suggest that hip actuation is preferred when the heel strike is advanced in which the trailing leg still can exert push-off impulse but delayed. Our simulation indicates that as the delayed push-off is still exerted along the trailing leg, a portion of it acts perpendicular to the new COM motion trajectory after the stance leg switch. Therefore, it must be wasted (converted to heat) [21] stretching the stance leg. In human walking, such stretch might also cancel a significant portion of mechanical energy stored in ligaments and tendons during the heel strike which is usually released in the subsequent push-off [22–24]. The higher than muscle active work performance walking efficiencies observed during walking indicates the share of elasticity in walking [20,25].

A tradeoff in mechanical workload between ankle extensors during push-off and hip flexors during terminal stance and early swing, in addition to contralateral hip extensors during early to midstance is noted [26]. It suggests that delayed push-off may impose a significant mechanical workload on the ankle extensors, potentially making it costlier in terms of energy expenditure compared to hip actuation. Furthermore, Hu et al. [27] have shown that the mechanical energy generated by the hip increases significantly during push-off, especially at faster walking speeds, implying that the hip muscles contribute to propel the body forward by increasing angular motion. This proposes that hip actuation could be a more efficient mechanism compared to delayed push-off in terms of mechanical or energetic cost [28].

In conclusion, based on simulation and a powered simple walking model we could demonstrate that the maximum possible push-off exertion bounds the step-up amplitude height or the minimum walking speed for the preferred approach to energize a gait. For cases when the pre-emptive push-off is sub-optimal, we compared the mechanical implications of actuating hip versus ankle. We could show that delayed push-off exertion entails stretching the stance leg which must waste a portion of the exerted mechanical energy to heat. Such a stretch perhaps also cancels a portion of the mechanical energy stored during the heel strike that contributes to the following step transition. Investigating prior research, we were also able to identify indications that humans also may prefer the hip actuation to delayed push-offs. In addition to leg swing frequency energetic cost [10,29] that is suggested to limit the lower of the step length, the delayed push-off cost may also be another determinant of the minimum step length based on the mechanical work. A shorter than preferred step length advances the heel strike that limits the pre-emptive push-off generation. Hence, not only the step energy dissipation elevates, but also the mechanical cost of delayed push-off must interrupt its performance. Nevertheless, the existence of the delayed push-off even in smooth nominal walking might indicate that humans’ sensory system must provide the required information which requires a processing time to substitute delayed push-off with the stance leg’s hip actuation.

## References

[1] A.D. Kuo, A Simple Model of Bipedal Walking Predicts the Preferred Speed–Step Length Relationship, Journal of Biomechanical Engineering 123 (2001) 264–269. 10.1115/1.1372322.

[2] J.E.A. Bertram, Constrained optimization in human walking: cost minimization and gait plasticity, Journal of Experimental Biology 208 (2005) 979–991. 10.1242/jeb.01498.

[3] A.D. Kuo, Energetics of actively powered locomotion using the simplest walking model, J Biomech Eng 124 (2002) 113–120. 10.1115/1.1427703.

[4] S.M. O’Connor, H.Z. Xu, A.D. Kuo, Energetic cost of walking with increased step variability, Gait & Posture 36 (2012) 102–107. 10.1016/j.gaitpost.2012.01.014.

[5] O. Darici, H. Temeltas, A.D. Kuo, Optimal regulation of bipedal walking speed despite an unexpected bump in the road, PLoS ONE 13 (2018) e0204205. 10.1371/journal.pone.0204205.

[6] A.D. Kuo, J.M. Donelan, A. Ruina, Energetic consequences of walking like an inverted pendulum: step-to-step transitions, Exerc Sport Sci Rev 33 (2005) 88–97. 10.1097/00003677-200504000-00006.

[7] N.F.J. Waterval, M.-A. Brehm, H.E. Ploeger, F. Nollet, J. Harlaar, Compensations in lower limb joint work during walking in response to unilateral calf muscle weakness, Gait & Posture 66 (2018) 38– 44. 10.1016/j.gaitpost.2018.08.016.

[8] K.E. Zelik, T.-W.P. Huang, P.G. Adamczyk, A.D. Kuo, The role of series ankle elasticity in bipedal walking, Journal of Theoretical Biology 346 (2014) 75–85. 10.1016/j.jtbi.2013.12.014.

[9] J.M. Donelan, R. Kram, A.D. Kuo, Mechanical work for step-to-step transitions is a major determinant of the metabolic cost of human walking, Journal of Experimental Biology 205 (2002) 3717–3727. 10.1242/jeb.205.23.3717.

[10] J. Doke, A.D. Kuo, Energetic cost of producing cyclic muscle force, rather than work, to swing the human leg, Journal of Experimental Biology 210 (2007) 2390–2398. 10.1242/jeb.02782.

[11] J.E.A. Bertram, S.J. Hasaneini, Neglected losses and key costs: tracking the energetics of walking and running, J Exp Biol 216 (2013) 933–938. 10.1242/jeb.078543.

[12] J.M. Caputo, S.H. Collins, Prosthetic ankle push-off work reduces metabolic rate but not collision work in non-amputee walking, Sci Rep 4 (2014) 7213. 10.1038/srep07213.

[13] O. Darici, H. Temeltas, A.D. Kuo, Anticipatory Control of Momentum for Bipedal Walking on Uneven Terrain, Sci Rep 10 (2020) 540. 10.1038/s41598-019-57156-6.

[14] O. Darici, A.D. Kuo, Humans plan for the near future to walk economically on uneven terrain, Proc. Natl. Acad. Sci. U.S.A. 120 (2023) e2211405120. 10.1073/pnas.2211405120.

[15] O. Darici, A.D. Kuo, Humans optimally anticipate and compensate for an uneven step during walking, eLife 11 (2022) e65402. 10.7554/eLife.65402.

[16] J.S. Matthis, S.L. Barton, B.R. Fajen, The critical phase for visual control of human walking over complex terrain, Proc. Natl. Acad. Sci. U.S.A. 114 (2017). 10.1073/pnas.1611699114.

[17] J.S. Matthis, J.L. Yates, M.M. Hayhoe, Gaze and the Control of Foot Placement When Walking in Natural Terrain, Curr Biol 28 (2018) 1224-1233.e5. 10.1016/j.cub.2018.03.008.

[18] S.J. Hasaneini, J.E.A. Bertram, C.J.B. Macnab, Energy-optimal relative timing of stance-leg push-off and swing-leg retraction in walking, Robotica 35 (2017) 654–686. 10.1017/S0263574715000764.

[19] P.G. Adamczyk, A.D. Kuo, Redirection of center-of-mass velocity during the step-to-step transition of human walking, J Exp Biol 212 (2009) 2668–2678. 10.1242/jeb.027581.

[20] A.S. Voloshina, A.D. Kuo, M.A. Daley, D.P. Ferris, Biomechanics and energetics of walking on uneven terrain, Journal of Experimental Biology (2013) jeb.081711. 10.1242/jeb.081711.

[21] R.McN. Alexander, Energy-Saving Mechanisms in Walking and Running, Journal of Experimental Biology 160 (1991) 55–69. 10.1242/jeb.160.1.55.

[22] N. Papachatzis, D.R. Slivka, I.I. Pipinos, K.K. Schmid, K.Z. Takahashi, Does the Heel’s Dissipative Energetic Behavior Affect Its Thermodynamic Responses During Walking?, Front. Bioeng. Biotechnol. 10 (2022) 908725. 10.3389/fbioe.2022.908725.

[23] R. Riemer, A. Shapiro, Biomechanical energy harvesting from human motion: theory, state of the art, design guidelines, and future directions, J NeuroEngineering Rehabil 8 (2011) 22. 10.1186/1743-0003-8-22.

[24] J. Ros, J.M. Font-Llagunes, A. Plaza, J. Kövecses, Dynamic considerations of heel-strike impact in human gait, Multibody Syst Dyn 35 (2015) 215–232. 10.1007/s11044-015-9460-0.

[25] G.A. Cavagna, N.C. Heglund, C.R. Taylor, Mechanical work in terrestrial locomotion: two basic mechanisms for minimizing energy expenditure, Am J Physiol 233 (1977) R243–261. 10.1152/ajpregu.1977.233.5.R243.

[26] S.N. Fickey, M.G. Browne, J.R. Franz, Biomechanical effects of augmented ankle power output during human walking, Journal of Experimental Biology (2018) jeb.182113. 10.1242/jeb.182113.

[27] Z. Hu, L. Ren, D. Hu, Y. Gao, G. Wei, Z. Qian, K. Wang, Speed-Related Energy Flow and Joint Function Change During Human Walking, Front. Bioeng. Biotechnol. 9 (2021) 666428. 10.3389/fbioe.2021.666428.

[28] Y. Ohta, H. Yano, R. Suzuki, M. Yoshida, N. Kawashima, K. Nakazawa, A two-degree-of-freedom motor-powered gait orthosis for spinal cord injury patients, Proc Inst Mech Eng H 221 (2007) 629– 639. 10.1243/09544119JEIM55.

[29] J. Doke, J.M. Donelan, A.D. Kuo, Mechanics and energetics of swinging the human leg, Journal of Experimental Biology 208 (2005) 439–445. 10.1242/jeb.01408.

